# PKA drives an increase in AMPA receptor unitary conductance during LTP in the hippocampus

**DOI:** 10.1101/2020.06.08.136762

**Authors:** Pojeong Park, John Georgiou, Thomas M. Sanderson, Kwang-Hee Ko, Heather Kang, Ji-il Kim, Clarrisa A. Bradley, Zuner A. Bortolotto, Min Zhuo, Bong-Kiun Kaang, Graham L. Collingridge

## Abstract

Long-term potentiation (LTP) at hippocampal CA1 synapses can be expressed by an increase either in the number (N) of AMPA (α-amino-3-hydroxy-5-methyl-4-isoxazole propionic acid) receptors or in their single channel conductance (γ). Here we have established how these distinct synaptic processes contribute to the expression of LTP in hippocampal slices obtained from young adult rodents. LTP induced by compressed theta burst stimulation (TBS), with a 10 s inter-episode interval, involved purely an increase in N (LTP_N_). In contrast, either a spaced TBS, with a 10 min inter-episode interval, or a single TBS, delivered when PKA was activated, resulted in LTP that was associated with a transient increase in γ (LTP_γ_). This γ increase was due to the insertion of calcium-permeable (CP)-AMPA receptors. Activation of CaMKII was necessary and sufficient for LTP_N_ whilst PKA was additionally required for LTP_γ_. Thus, two mechanistically distinct forms of LTP co-exist at these synapses.

## INTRODUCTION

Long-term potentiation (LTP) of synaptic function is considered the major process underlying learning and memory ^1^ where it is involved in synaptic engram formation ^2,3^. yet the underlying cellular mechanisms remain incompletely understood. The best-characterized form of LTP occurs at the Schaffer collateral-commissural pathway (SCCP) in the hippocampus, where it is triggered by synaptic activation of NMDA (N-methyl-D-aspartate) receptors ^4^ and is expressed as a persistent increase in AMPA (α-amino-3-hydroxy-5-methyl-4-isoxazole propionic acid) receptor-mediated synaptic transmission ^5^. This modification is primarily due to a functional modulation of AMPA receptors (AMPARs), which may involve a change in the number of active channels (N) (termed LTP_N_) and/or their single-channel conductance (γ) properties (termed LTPγ) (e.g., ^6–9^). Whilst there is considerable evidence that LTP_N_ is triggered by activation of Ca^2+^/calmodulin-dependent kinase II (CaMKII) ^10,11^ and involves exocytosis and lateral diffusion of AMPARs ^12,13^, the mechanisms underlying LTP_γ_ are largely unknown. The two most likely molecular mechanisms involve (i) CaMKII-mediated phosphorylation of Ser831 of GluA1, which can result in an increase in the time AMPARs dwell in higher conductance states ^14–16^ or (ii) the insertion of calcium-permeable AMPA receptors (CP-AMPARs), which have a higher γ than their calcium-impermeable (CI) counterparts ^17,18^.

In the present study, we have tested the hypothesis that LTP_γ_ is due to the insertion of CP-AMPARs in young adult rodents using two theta burst stimulation (TBS) induction protocols that differed only in the timing between episodes, and applied peak-scaled non-stationary fluctuation analysis (NSFA) ^19–21^ to estimate γ before and after the induction of LTP ^6,15,22–25^. We found that the compressed TBS protocol (cTBS-inter-episode interval of 10 s) resulted exclusively in LTP_N_, for which CaMKII was both necessary and sufficient. In contrast, a spaced TBS protocol (sTBS - inter-episode interval of 10 min) resulted in a transient increase in γ, lasting ~15 min, which was due to the insertion of CP-AMPARs and required both CaMKII and PKA. Insertion of CP-AMPARs mediates both the initial expression of LTP_γ_, by enhancing the net synaptic unitary conductance, and helps trigger the processes that lead to a persistent increase in synaptic efficacy that outlasts the increase in γ. Since the PKA-dependent form of LTP also requires *de novo* protein synthesis and has stimulation features similar to spaced behavioural learning, LTP_γ_ is likely to underlie the formation of synaptic engrams and long-term memory.

## RESULTS

### An increase in γ is specifically triggered by a sTBS protocol

Simultaneous field excitatory postsynaptic potential (fEPSP) recordings from stratum radiatum and somatic whole-cell recordings were obtained in response to baseline stimulation of two independent SCCP inputs (Fig. 1a). TBS was delivered to one input (test), while the second input served as a control for stability and heterosynaptic effects (Fig. 1c, d). Synaptic potentiation was quantified and γ was estimated using NSFA (Fig. 1e, f), as described previously ^6^. To optimise the estimates of γ we used minimal stimulation and restricted our measurements to the first 20-30 min following TBS, since γ estimates are extremely sensitive to small fluctuations in series resistance ^20^. Thus, our study focused on the induction and initial expression mechanisms of LTP.

**Fig. 1.**
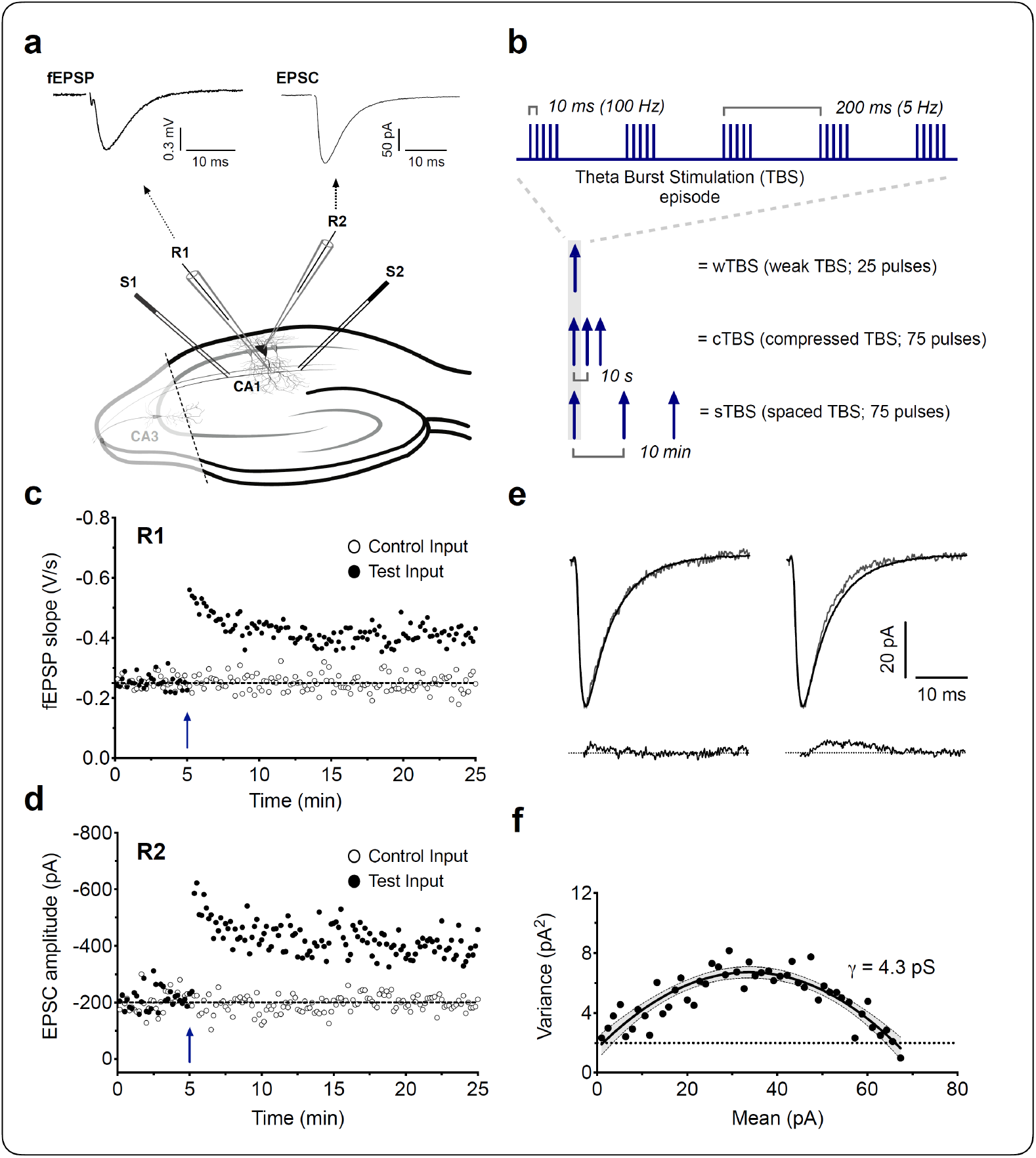
LTP and non-stationary fluctuation analysis (NSFA) methodology. **a**, Schematic of a hippocampal brain slice for LTP experiments, and the positioning of recording (R1, R2) and stimulating (S1, S2) electrodes. The CA3 region was cut (dashed line) to reduce neuronal excitability. Representative field and whole cell responses (fEPSP and EPSC), simultaneously obtained from CA1 neurons. Five consecutive responses were averaged and the stimulus artifacts were blanked for clarity. **b,** Induction protocols for weak, compressed and spaced TBS (wTBS, cTBS and sTBS) are graphically summarized. **c**, Representative fEPSP recordings for LTP evoked by a single episode of TBS (weak TBS, blue arrow). **d**, Simultaneously obtained EPSC recordings. **e**, Upper traces are two sets of representative waveforms for individual sweeps (thin lines), superimposed with the scaled mean of 57 EPSCs (thick lines). Lower traces are the subtraction of the scaled mean from the representative individual EPSCs. **f**, Corresponding current-variance relationship to estimate the unitary conductance (γ). Fluctuation of the individual decays was plotted against the mean EPSC. Solid line is a parabolic fit with 95% confidence intervals (shaded). Dotted line, the background average variance.

In the first series of experiments we delivered three episodes of TBS, with each episode comprising 5 shocks at 100 Hz delivered 5 times at 5 Hz (i.e., 75 stimuli in total; see Fig. 1b schematic); in interleaved experiments we either delivered these three episodes as a cTBS (10 s inter-episode interval) or as a sTBS (10 min inter-episode interval). We referred to the resultant potentiation as cLTP (Fig. 2a-i) and sLTP (Fig. 2j-r), respectively. In response to cTBS there was a substantial cLTP (Fig. 2a), with EPSC amplitudes increasing to 212 ± 11% of baseline, averaged over the first 10 min after induction (Fig. 2b). For 22 neurons from 15 rats (n = 22/15), we obtained γ estimates in 10 min epochs and found it to be unaltered throughout (Fig. 2c-g). The γ values were 5.1 ± 0.3 pS, (baseline), 5.3 ± 0.4 pS (first 10 min epoch post cTBS; LTP_10’_; t_21_ = 1.23, *p* = 0.2327, *vs* baseline, paired Student’s *t*-test) and 5.2 ± 0.4 pS (second 10 min epoch post cTBS; LTP_20’_; t_21_ = 0.33, *p* = 0.7452; Fig. 2d). The control input was also stable throughout (4.9 ± 0.4 pS, 4.5 ± 0.3 pS and 4.8 ± 0.3 pS at the corresponding time-points; Fig. 2d). The lack of change in γ was also clearly evident in the plots from individual experiments for control (Fig. 2e) and test inputs (Fig. 2f) and in the cumulative distribution plots (Fig. 2g). The lack of change in γ was observed over a wide range of cLTP magnitudes (Fig. 2h).

**Fig. 2.**
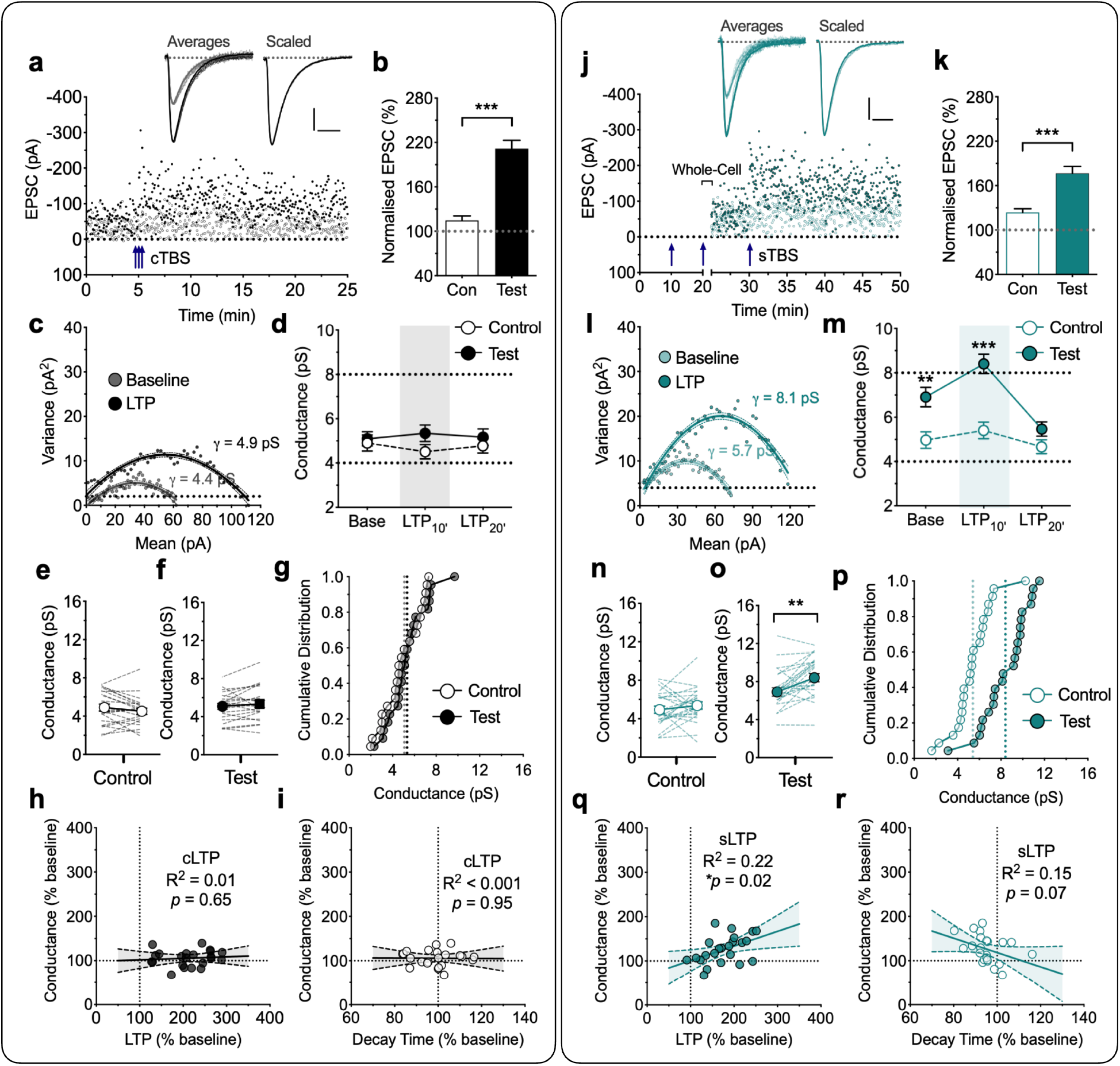
Increased AMPA receptor unitary conductance (γ) during sLTP, but not cLTP. **a**, A representative LTP experiment with sample traces for baseline and post TBS – the mean of selected records for analysis, superimposed with peak-scaled individual traces (10 successive sweeps, thin lines; baseline = grey, LTP = black). Scaled trace is from the baseline normalized to the LTP. Scale bars: 20 pA and 10 ms. Two inputs were stimulated alternately and cTBS (3 x TBS with an inter-episode interval of 10 sec; blue arrows) delivered to one input (filled symbols) with the second input (open symbols) serving as a control (Con). **b**, Levels of cTBS-induced LTP (cLTP) for control and test inputs, quantified during the 10 min epoch after the induction (mean ± SEM, n = 22 neurons from 15 animals; t_21_ = 8.55, \*\*\**p* < 0.001, paired Student’s *t*-test). **c**, Corresponding current-variance relationship of the EPSCs for the test input. The unitary channel conductance (γ) of AMPA receptors was estimated during baseline (grey) and after the induction of LTP (LTP_10’_; black). **d**, Grouped comparison of control and test input γ estimates for baseline and the initial 10 min epoch (LTP_10’_) and the subsequent 10 min epoch (LTP_20’_). **e-f**, Summary plot for the γ at baseline (left) and LTP_10’_ (right) for control (**e**) and test (**f**) inputs. Individual values (circles) from each neuron are connected by lines. **g**, Cumulative distribution of the same data set for LTP_10’_. Dotted lines indicate the mean values for each input. **h-i**, Analysis of the relationships of γ with, LTP (**h**) and EPSC decay time (**i**). Linear regression with 95% confidence intervals (shaded) for the amount of cLTP and the corresponding level of γ. **j-p**, Equivalent analysis for the LTP induced by sTBS (3 x TBS at inter-episode interval of 10 min; see arrows). The whole-cell recordings were obtained after the second TBS. This was necessary due to the lability of LTP washout. **m-o**, statistical analysis between control and test pathways (**m**) and within pathway analysis for control (**n**) and test (**o**) pathway reveals a time- and pathway-dependent increase in γ. Note that higher conductance was observed in the test input (**o**) compared to the control (**n**) under the “baseline” state, suggesting that the first + second TBS were sufficient to increase γ. The third TBS triggered a small but discernible further increase in γ (n = 23/17, t_22_ = 3.75, \*\**p* < 0.01, paired Student’s t-test). **q-r**, Analysis of the relationships of γ with LTP (**q**) and decay time of EPSCs (**r**).

In response to sTBS the results were strikingly different. For this set of experiments, whole-cell recordings were obtained shortly after delivery of the second TBS episode and the effects of the third TBS was evaluated (Fig. 2j). This method was necessary because of the rapid wash-out of LTP with low access whole-cell recordings. In response to the third TBS there was a substantial additional LTP, with EPSC amplitudes increasing to 177 ± 9% of baseline, averaged over the first 10 min after induction (Fig. 2k). The estimate of γ upon break in was significantly higher (6.9 ± 0.4 pS) compared to the control input (4.9 ± 0.4 pS; Fig. 2n-o; t_22_ = 3.22, *p* = 0.0039, paired Student’s *t*-test) and this was further increased in response to the third episode of TBS to 8.4 ± 0.4 pS (LTP_10’_; t_22_ = 3.75, *p* = 0.0011, Fig. 2l, m, o, p; n = 23/17). However, when we quantified γ at 10-20 minutes after the last TBS, the value (5.5 ± 0.3 pS) was no longer significantly different from the control input (LTP_20’_; t_22_ = 2.01, *p* = 0.0570, paired Student’s *t*-test; Fig. 2m). In contrast to the test input, sTBS did not result in a significant γ change in the control input (4.9 ± 0.4 pS, 5.4 ± 0.4 pS and 4.6 ± 0.3 pS at the corresponding time points; Fig. 2m-n). Thus, the increase in γ is specifically related to sLTP. Furthermore, this increase in γ positively correlated with the magnitude of sLTP (Fig. 2q).

Since sLTP, but not cLTP, is associated with the insertion of CP-AMPARs ^26,27^ these results suggest that CP-AMPARs may account for the increase in γ. CP-AMPARs have slightly faster decay kinetics (τdecay) than CI-AMPARs ^25,28^, which can be detected using single exponential fits to EPSC decays. We found that cLTP was not associated with an alteration in τdecay (Fig. 2i, Table 1; t_21_ = 0.66, *p* = 0.5146, paired Student’s *t*-test), whereas sLTP was associated with a highly significant decrease in τ_decay_ (Table 1; *p* = 0.0051, t_22_ = 3.11, paired Student’s *t*-test). A regression analysis showed a trend for the τ_decay_ to be inversely related with the increase in γ (Fig. 2r; *p* = 0.0712, F_(1, 21)_ = 3.61). Therefore, the kinetic analysis provides additional support for the notion that insertion of CP-AMPARs occurs during the induction of LTP in response to a sTBS.

**Table 1.** Summary of EPSC properties for the various experimental protocols and conditions.

| Protocol | $\gamma$ (pS) | | EPSC (%) | $\tau_{\text{rise}}$ (ms) | | $\tau_{\text{decay}}$ (ms) | | N |
| --- | --- | --- | --- | --- | --- | --- | --- | --- |
| Compressed TBS | $5.09 \pm 0.32$ | $5.34 \pm 0.37$ | $212 \pm 11^{***}$ | $1.23 \pm 0.06$ | $1.20 \pm 0.07$ | $7.17 \pm 0.19$ | $7.07 \pm 0.23$ | 22/15 |
| Spaced TBS | $6.91 \pm 0.44$ | $8.40 \pm 0.44^{**}$ | $177 \pm 9^{***}$ | $1.16 \pm 0.06$ | $1.15 \pm 0.05$ | $7.09 \pm 0.19$ | $6.75 \pm 0.20^{**}$ | 23/17 |
| wTBS with rolipram | $4.86 \pm 0.43$ | $8.02 \pm 0.58^{***}$ | $234 \pm 14^{***}$ | $1.22 \pm 0.06$ | $1.16 \pm 0.05$ | $7.19 \pm 0.20$ | $6.68 \pm 0.15^{***}$ | 21/15 |
| wTBS with PKA $C\alpha$ | $5.15 \pm 0.51$ | $7.79 \pm 0.80^{***}$ | $276 \pm 19^{***}$ | $1.02 \pm 0.06$ | $1.02 \pm 0.04$ | $6.75 \pm 0.22$ | $6.41 \pm 0.23^*$ | 17/13 |
| wTBS with PKA $C\alpha$ + IEM | $4.29 \pm 0.51$ | $4.32 \pm 0.54$ | $202 \pm 16^{***}$ | $1.00 \pm 0.07$ | $0.99 \pm 0.07$ | $6.73 \pm 0.25$ | $6.70 \pm 0.26$ | 16/13 |
| HI-CaMKII | $4.57 \pm .053$ | $4.57 \pm 0.52$ | $112 \pm 7$ | $1.01 \pm 0.06$ | $1.01 \pm 0.07$ | $6.99 \pm 0.34$ | $6.96 \pm 0.25$ | 14/11 |
| CaMKII | $4.71 \pm 0.57$ | $4.32 \pm .046$ | $169 \pm 15^{***}$ | $0.95 \pm 0.05$ | $0.93 \pm 0.04$ | $6.82 \pm 0.30$ | $6.75 \pm 0.29$ | 15/12 |
| CaMKII + PKA $C\alpha$ | $4.64 \pm 0.39$ | $6.48 \pm 0.42^{***}$ | $178 \pm 10^{***}$ | $1.05 \pm 0.07$ | $1.03 \pm 0.08$ | $7.06 \pm 0.26$ | $6.69 \pm 0.19^*$ | 18/15 |
| CaMKII + PKA $C\alpha$ + IEM | $4.26 \pm 0.33$ | $4.13 \pm 0.40$ | $128 \pm 9^*$ | $1.01 \pm 0.04$ | $1.00 \pm 0.04$ | $6.91 \pm 0.20$ | $6.87 \pm 0.22$ | 20/15 |
| HI-CaMKII + PKA $C\alpha$ | $4.24 \pm 0.35$ | $4.22 \pm 0.40$ | $112 \pm 9$ | $1.01 \pm 0.03$ | $0.97 \pm 0.03$ | $7.07 \pm 0.27$ | $7.05 \pm 0.26$ | 16/14 |
| CaMKII + IEM | $4.07 \pm 0.39$ | $4.04 \pm 0.66$ | $165 \pm 17^{**}$ | $0.97 \pm 0.03$ | $0.98 \pm 0.03$ | $6.75 \pm 0.22$ | $6.78 \pm 0.34$ | 8/8 |

\* $p < 0.05$ ; \*\* $p < 0.01$ ; \*\*\* $p < 0.001$ for baseline vs. LTP in paired Student's *t*-test.

### The role of PKA in LTP_γ_

It is established that elevating cAMP by, for example, use of the phosphodiesterase 4 inhibitor rolipram, enables a weak stimulus to generate an enhanced PKA-dependent form of LTP ^29^. Previously, we found that in the presence of rolipram a weak TBS, comprising one episode of TBS, generated an LTP that is largely dependent on the insertion of CP-AMPARs ^26^. Here we used this same method as an independent means to investigate whether insertion of CP-AMPARs are responsible for the increase in γ. Since only one TBS is required to induce the PKA-dependent form of LTP in the presence of rolipram we could make γ measurements before and after the full induction of LTP. As illustrated in Fig. 3a-b, a single episode of TBS (wTBS; comprising 25 stimuli), when delivered in the presence of rolipram (1 μM), generated a robust LTP (234 ± 14 % of baseline for test vs. 121 ± 6 % for control input). We found that this LTP was also associated with a transient increase in γ (baseline = 4.9 ± 0.4 pS, LTP_10’_ = 8.0 ± 0.6 pS; t_20_ = 5.90, *p* < 0.0001) that returned to baseline by the second 10 min epoch (LTP_20’_ = 5.4 ± 0.3 pS; t_20_ = 1.39, *p* = 0.1810) following the wTBS (n = 21/15; Fig. 3c, d, f, g). This potentiation required the wTBS since the control input was largely unaffected (5.1 ± 0.3 pS, 5.4 ± 0.5 pS and 4.8 ± 0.3 pS at the corresponding time points; Fig. 3d, e) and since the baseline γ values in the presence of rolipram were not significantly different to the baseline γ values in its absence (Fig. 3d-f; Table 1). As was the case with the sLTP, the size of the change in γ correlated with the magnitude of LTP (*p* = 0.0024, F_(1, 19)_ = 12.27; Fig. 3h). Additionally, there was an associated reduction in τ_decay_ (*p* = 0.0007, t_20_ = 3.99, paired Student’s *t*-test; Table 1) that also negatively correlated with the increased γ (*p* = 0.0199, F_(1,19)_ = 6.46; Fig. 3i). These results further support the idea that insertion of CP-AMPARs mediates LTP_γ_.

**Fig. 3.**
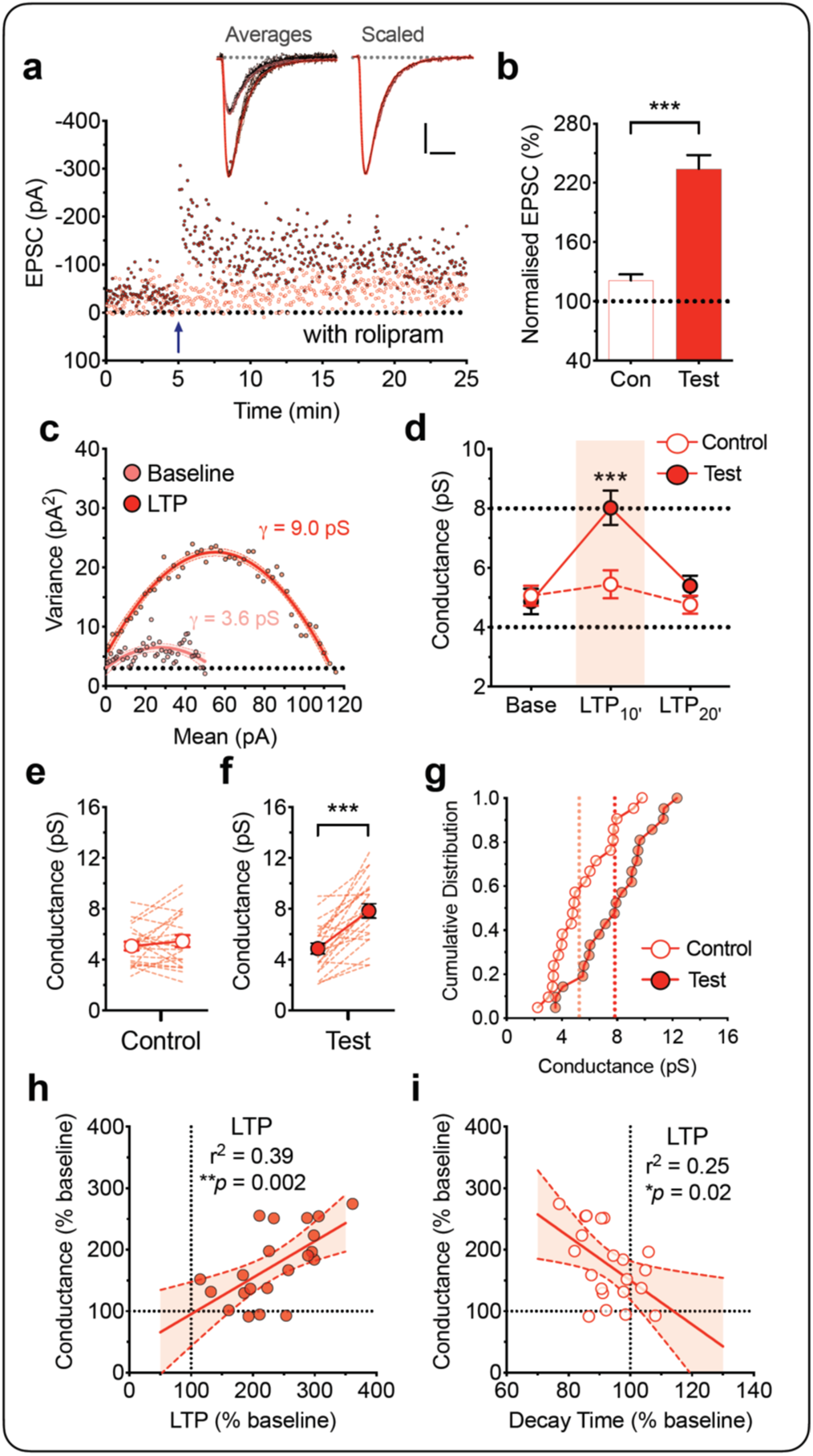
Increased AMPA receptor unitary conductance (γ) during LTP in the presence of rolipram. Equivalent experiments to those illustrated in Fig. 2 for the LTP induced by a wTBS (a single episode of TBS) in the presence of rolipram (1 μM; n = 21/15).

To more specifically test the requirement of PKA for driving alterations in γ, we included the catalytic subunit of PKA (PKA Cα; 300 U/mL) in the patch solution (Fig. 4). This treatment had little effect on the control input (Fig. 4a). However, as was the case with rolipram, the wTBS in the presence of PKA Cα generated a robust potentiation (Fig. 4a) that was associated with an increase in γ (Fig. 4b, c). The levels quantified during baseline and 10 min post TBS (LTP_10’_) were 5.2 ± 0.5 pS and 7.8 ± 0.8 pS (t_16_ = 5.80,*p* < 0.0001, paired Student’s *t*-test; n = 17/13; Fig. 4b). Once again, the increase in γ was only transient, since estimates of γ made between 10 and 20 min following the wTBS (i.e. LTP_20’_) were not significantly different from baseline (5.3 ± 0.5 pS; t_16_ = 0.37, *p* = 0.7163, paired Student’s *t*-test; Fig. 4b).

**Fig. 4.**
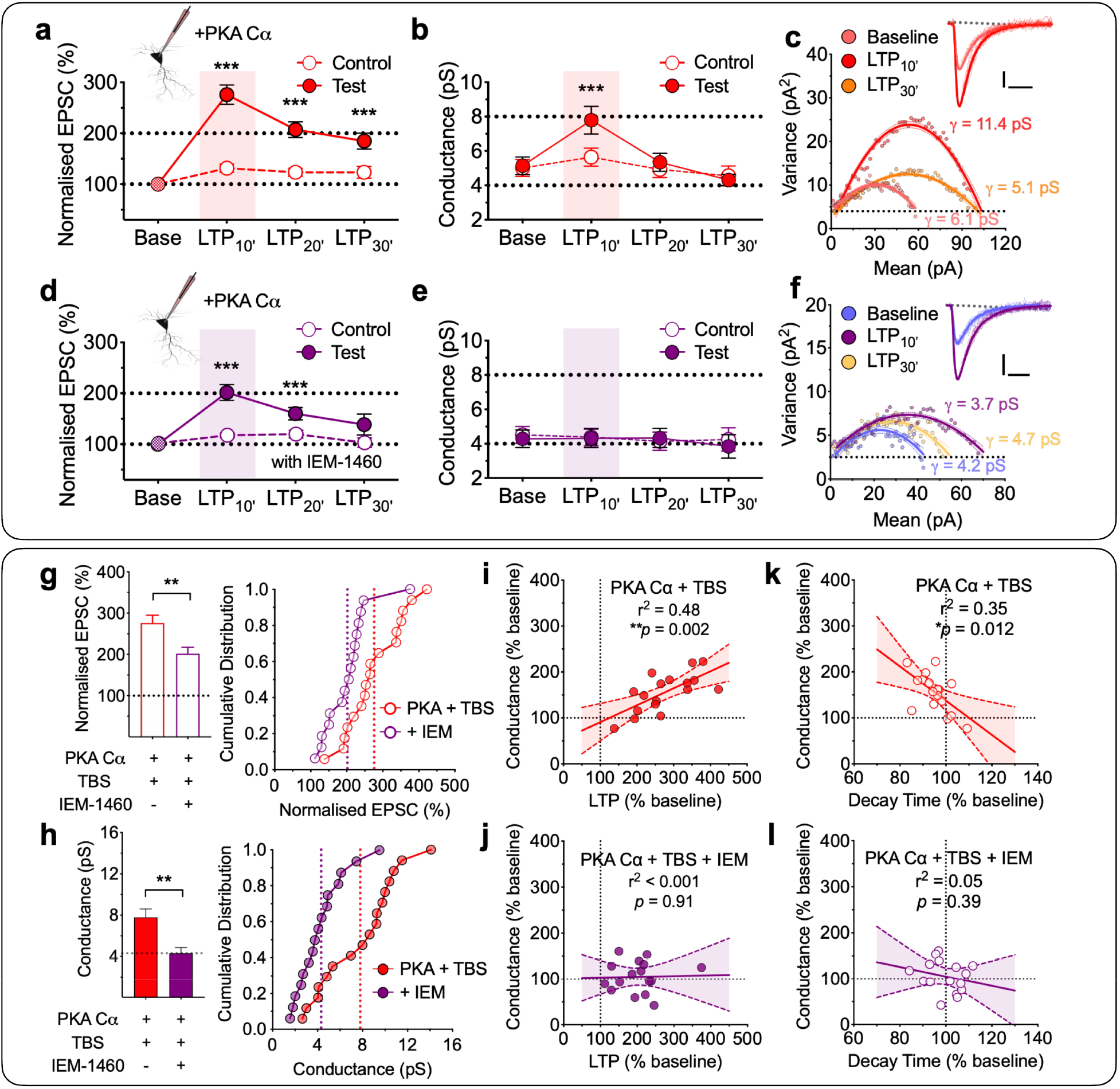
wTBS with PKA Cα transiently increases γ via CP-AMPAR insertion. **a-b**, a wTBS in the presence of intracellular PKA Cα (300 U/mL) transiently increased γ (n = 17/13, t_16_ = 5.80, paired Student’s t-test, \*\*\**p* < 0.001). EPSCs (**a**) and γ (**b**) were analyzed in 10-min bins. A single episode of TBS (at time marked by an arrow) was delivered to one input (filled symbols) with the second input (open symbols) serving as a control; base = baseline. **c**, A representative current-variance plot for PKA Cα plus wTBS for baseline, the first 10 min (LTP_10’_) and the last 10 min of LTP (LTP30’). Sample traces were obtained from baseline and LTP_10’_. Scale bars: 10 pA and 10 ms. **d-f**, Equivalent experiments in the presence of IEM-1460 (IEM, 30 μM; n = 16/13). **g-h**, Quantification of the levels of LTP (**g**) and γ (**h**) measured during the 10 min after wTBS with cumulative distributions (right). **i-l**, Analysis of the relationships between γ and LTP (**i** and **j**) and EPSC decay time (**k** and **l**) for PKA Cα + wTBS (**i** and **k**) and PKA Cα + wTBS + IEM (**j** and **l**).

In interleaved experiments (Fig. 4d-f), we examined the effects of the CP-AMPAR blocker, IEM-1460 (30 μM, IEM). In the presence of bath applied IEM and PKA Cα in the patch pipette, the level of LTP triggered by the wTBS was significantly less than in its absence (202 ± 16% vs. 276 ± 19% of baseline, 10 min after wTBS; t_31_ = 3.01, *p* = 0.0052, unpaired Student’s *t*-test; Fig. 4d, g), consistent with a component of LTP being generated by the insertion of CP-AMPARs when PKA is activated ^26,30–32^. IEM completely prevented the transient increase in γ (baseline vs. LTP_10’_; 4.3 ± 0.5 pS vs. 4.3 ± 0.5 pS; t_15_ = 0.08, *p* = 0.9338, paired Student’s *t*-test; Fig. 4e, h; n = 16/13; also see Table 1). There was a strong correlation between the increase in γ with both the magnitude of LTP (Fig. 4i; *p* = 0.0021, F_(1,15)_ = 13.72) and the decrease in τ_decay_ (Fig. 4k; *p* = 0.0117, F_(1,15)_ = 8.24) when wTBS was delivered in the presence of PKA Cα, but there was no such correlations when IEM was also present (Fig. 4j, l).

In conclusion, we find that activation of PKA, that occurs during (i) a sTBS, (ii) a wTBS in the presence of rolipram, or (iii) a wTBS in the presence of the catalytic subunit of PKA, results in the transient insertion of CP-AMPARs and that these receptors are responsible for the increase in γ during the initial expression phase of LTP.

### The role of CaMKII in LTP_γ_

CaMKII has been demonstrated to be both necessary and sufficient for the induction of LTP ^10,11^. Consistent with this notion, when tested using a CaMKII selective antagonist, KN-62 (10 μM), we found that both cLTP and sLTP were substantially reduced (Fig. 5a, b). The levels of potentiation of 108 ± 8% (after 90 min of cTBS, n = 4 slices from individual animals; Fig. 5a) and 105 ± 7% (after 120 min of sTBS, n = 5 slices; Fig. 5b), respectively, were significantly less than that in the corresponding interleaved untreated control groups that potentiated 155 ± 5% (*p* = 0.0010, t_9_ = 4.79; unpaired Student’s *t*-test; n = 7 slices) and 159 ± 3% (*p* = 0.0001, t_13_ = 6.87; unpaired Student’s *t*-test; n = 10 slices), but were not significantly different from their respective control inputs (t_3_ = 1.50, *p* = 0.2315 and t_4_ = 0.66; *p* = 0.5426; paired Student’s *t*-test).

**Fig. 5.**
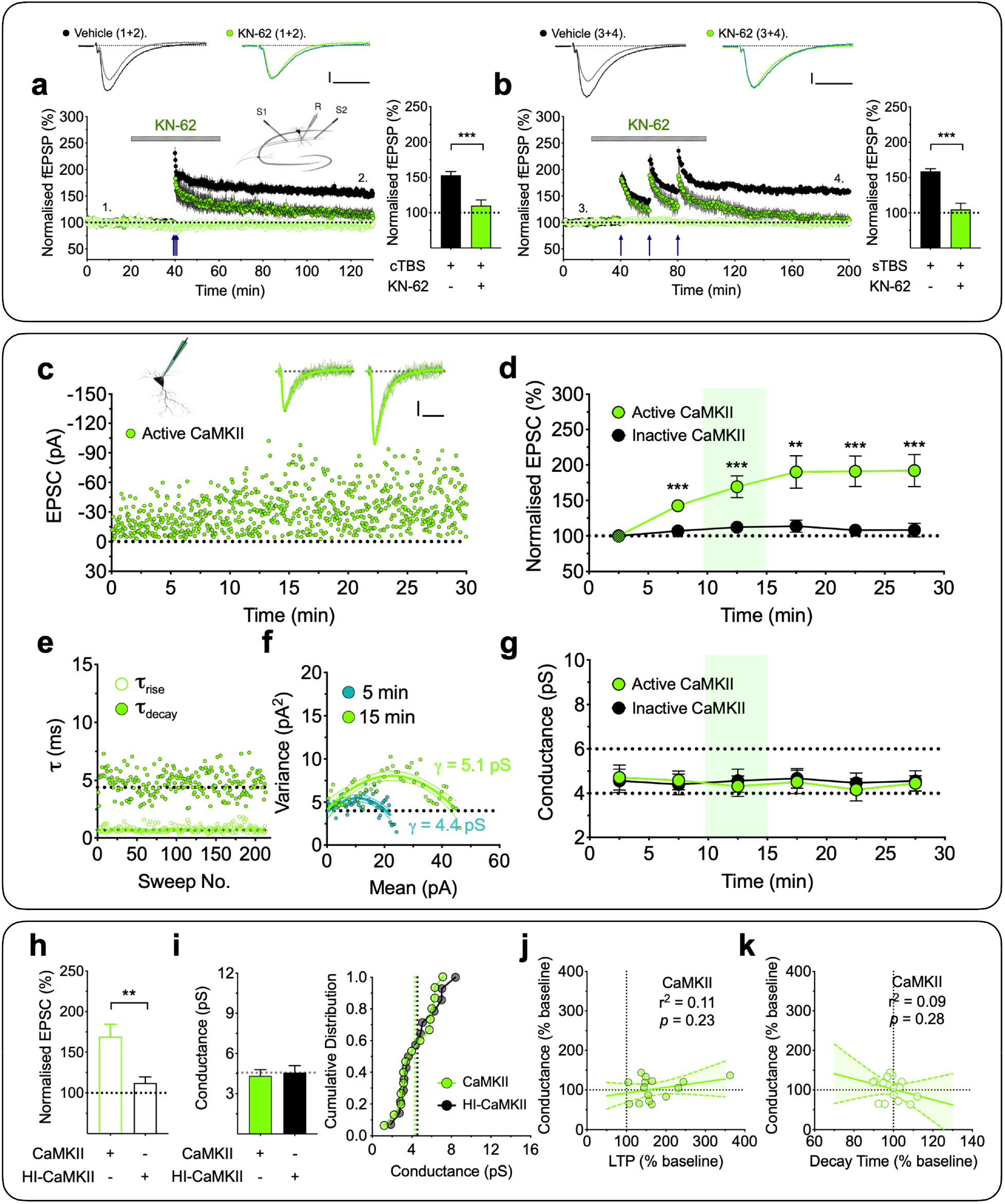
CaMKII does not affect γ. **a-b**, CaMKII-dependence of both forms of LTP. **a**, cLTP, measured using fEPSP recordings, was inhibited by the CaMKII inhibitor, KN-62 (10 μM; n = 4 animals; green). **b**, sLTP showed a similar sensitivity to KN-62 (n = 5). Interleaved control experiments (n = 7 and 10; black) are superimposed. The sample traces were obtained at the timing indicated by the numbers. Scale bars: 0.2 mV and 10 ms. **c**, A representative whole-cell recording with the inclusion of activated CaMKII (250 U/mL) in the internal solution. The sample traces are averages of selected records for analysis, superimposed with individual scaled traces (10 successive sweeps, thin lines) after 5 and 15 min of whole-cell recording. Scale bars: 10 pA and 10 ms. **d**, Pooled results (mean ± SEM, 5-min bins) for the effects on EPSC (%) by activated CaMKII (n = 15/12) and interleaved control, heat-inactivated, CaMKII (n = 14/11). **e**, Rise times (20-80%, τ_rise_) and decay time constants (τ_decay_) were plotted for the EPSCs used in the NSFA analysis for the neuron illustrated in **c**. **f**, Current-variance relationships for this neuron used to estimate γ over 5 min epochs starting at 0 and 10 min after commencing wholecell recording. **g**, Time course for the estimates of γ for active *vs.* inactive CaMKII. **h-i**, Quantification for the levels of LTP (**h**) and γ (**i**) measured over a 5 min epoch, commencing 10 min after starting whole-cell recording. **j-k**, Analysis of the relationships between γ and LTP (**j**) and EPSC decay time (**k**) for the active CaMKII experiments.

It has been suggested that the role of CaMKII in LTP involves an increase in γ ^14,15^. To further examine the role of CaMKII in LTP we interleaved experiments where we applied either active or inactive (heat inactivated) CaMKII (250 U/mL) via the patch pipette and delivered baseline (low frequency) stimulation to monitor basal synaptic transmission. Consistent with previous reports ^33,34^, activated CaMKII, but not inactive CaMKII, was sufficient to potentiate synaptic transmission (Fig. 5c, d, h). However, this potentiation was not associated with an increase in γ (Fig. 5f, g, i) or a change in rise and decay kinetics (Fig. 5e; see also Table 1). The respective γ values for baseline (i.e., first 5 min of recording) and 10-15 min of whole-cell recording were 4.7 ± 0.6 pS and 4.3 ± 0.5 pS (t_14_ = 0.74, *p* = 0.4740, paired Student’s *t*-test; n = 15/12; Fig. 5g). There was no correlation between γ change and either the magnitude of LTP (Fig. 5j; *p* = 0.2265, F_(1,13)_ = 1.61) or τ_decay_ (Fig. 5k; *p* = 0.2813, F_(1,13)_ = 1.26). We can conclude, therefore, that CaMKII alone can generate substantial potentiation that does not involve any alteration in γ.

### Activation of CaMKII and PKA are both necessary and sufficient for LTP_γ_

Since neither PKA alone nor CaMKII alone affected γ, we explored whether the combination of the two kinases may be sufficient for the effect. We, therefore, patch loaded PKA Cα (300 U/mL) with either the active or inactive forms of CaMKII (250 U/mL). In interleaved experiments, we found that PKA Cα + active CaMKII produced a robust potentiation of synaptic responses, specifically 178 ± 10% of baseline when quantified 15 min after whole-cell (Fig. 6a, b, f). In this case, the effect was also associated with an increase in γ (Fig. 6d, e, g). The levels of conductance for the baseline and potentiation (calculated between 10 to 15 min of recording) were 4.6 ± 0.4 pS and 6.5 ± 0.4 pS, respectively (t_17_ = 5.38, *p* = 0.0002, paired Student’s *t*-test; n = 18/15). Again, this effect was only transient, as the γ returned to baseline levels within 20-30 min of whole-cell recording (Fig. 6e). In contrast, inactive CaMKII plus PKA Cα, had no significant effect on synaptic transmission (112 ± 9%; Fig. 6b, f) or on γ (4.2 ± 0.4 pS vs. 4.2 ± 0.4 pS, n = 16/14; Fig. 6e, g). These results suggest that (i) both CaMKII and PKA are required for and (ii) their combined activity is sufficient for LTP_γ_ at these synapses.

**Fig. 6.**
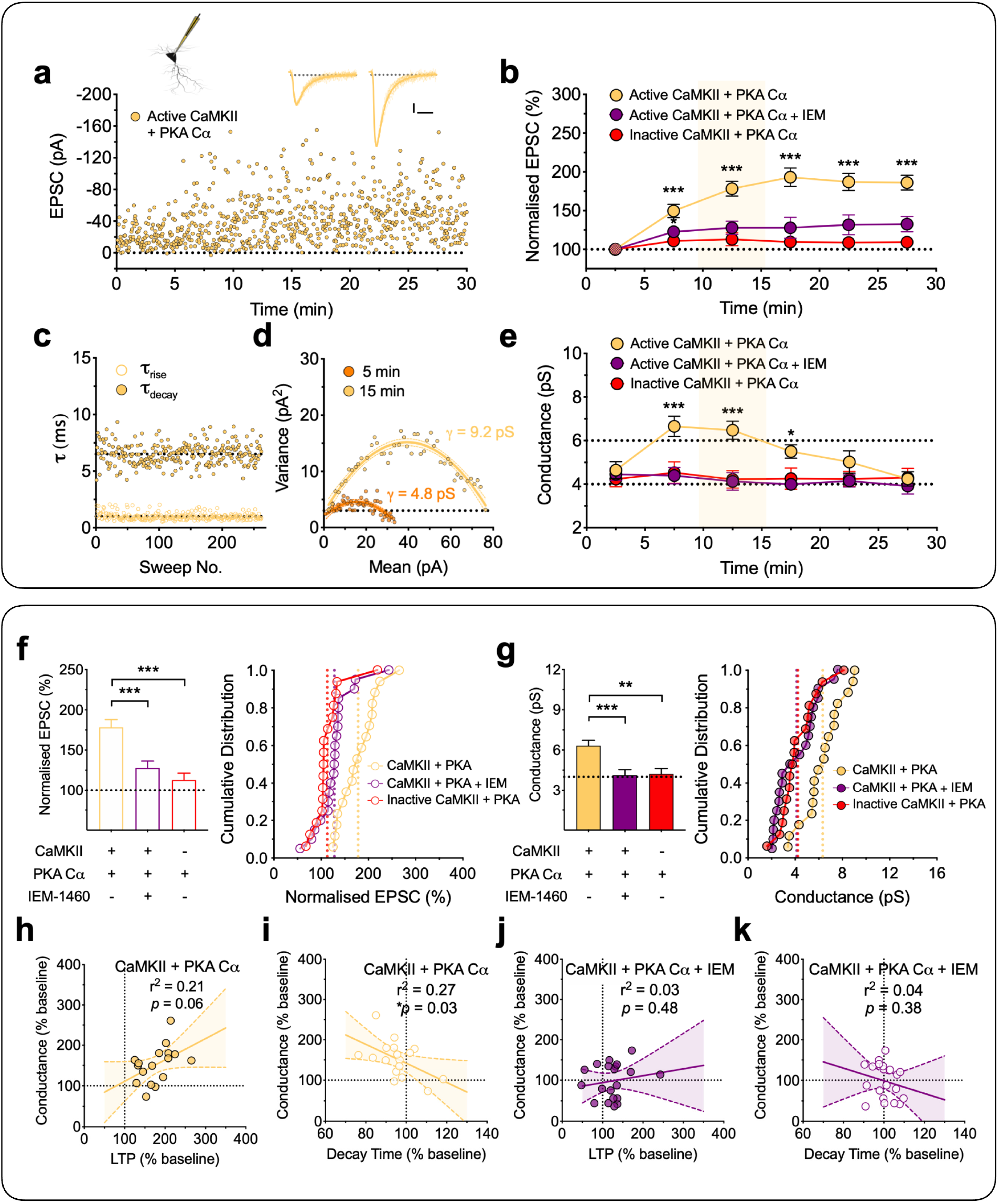
CaMKII plus PKA Cα results in a transient synaptic insertion of CP-AMPARs and increase in γ. Equivalent experiments as described in Fig. 5**c-k** but with the inclusion of activated CaMKII (250 U/mL) plus the catalytic subunit of PKA (PKA Cα, 300 U/mL) in the internal solution. Data were analyzed for active CaMKII + PKA Cα (n = 18/15), CaMKII + PKA Cα + IEM, 30 μM (n = 20/15) and heat-inactivated CaMKII + PKA Cα (n = 16/14). \**p* < 0.05, \*\**p* < 0.01 and \*\*\**p* < 0.001.

In additional interleaved experiments, the sensitivity to IEM was tested on the potentiation produced by CaMKII plus PKA Cα. Consistent with the involvement of CP-AMPARs, there was a reduced level of potentiation (Fig. 6b, f) and no change in γ in the presence of IEM (Fig. 6e, g). The respective amounts, quantified after 10-15 min of whole-cell recording, were 128 ± 9% of baseline (*p* = 0.0003 vs. CaMKII + PKA Cα, one-way ANOVA with Bonferroni’s correction) and 4.1 ± 0.4 pS (*p* = 0.0009 vs. CaMKII + PKA Cα, one-way ANOVA with Bonferroni’s correction; n = 20/15).

The correlations between the change in γ with the level of LTP and decay times for these experiments are summarized in figure 6h-k. There was a substantial correlation between the increase in γ and the level of potentiation (Fig. 6h) and the decrease in τ_decay_ (Fig. 6i) with CaMKII plus PKA Cα but no such correlation was found in the presence of IEM (Fig. 6j-k).

### The proportion of synaptically-incorporated CP-AMPARs during LTP_γ_

Together, the previous experiments provide multiple lines of evidence that LTP_γ_ is due to the insertion of CP-AMPARs into synapses that contain CI-AMPARs. In order to determine the relative proportions of each it was necessary to measure γ for synapses containing either just CI-AMPARs or just CP-AMPARs, under our recording conditions. To achieve this, we used lentivirus-driven CRISPR/Cas9 expression to delete GluA2 in a fraction of neurons *in vivo*, allowing a direct comparison between a knock-out (KO) and a wild-type (WT) neuron within each adult brain slice (Fig. 7a). When compared with uninfected neighbouring neurons, the KO cells showed a reduced AMPAR synaptic transmission (Fig. 7b) and an inwardly rectifying current-voltage relationship (Fig. 7c, d). The level of γ in KO neurons was significantly higher at 17.3 ± 1.2 pS (n = 16) compared to 4.6 ± 0.4 pS for WT neurons (n = 17; t_31_ = 10.09, *p* < 0.0001, unpaired Student’s *t*-test, data from 12 animals; Fig. 7e-h). This increase in γ in these neurons was associated with a decreased τ_decay_ from 6.8 ± 0.4 ms in WT to 5.3 ± 0.4 ms in KO neurons (t_31_ = 2.85, *p* = 0.0026, unpaired Student’s *t*-test, Fig. 7i). Assuming that these EPSCs were comprised of 100 and 0 % CP-AMPARs, respectively, then the increase in γ that we observed during sLTP can be explained by CP-AMPARs comprising ~30% of the synaptic current during the first 10 min following LTP induction.

**Fig. 7.**
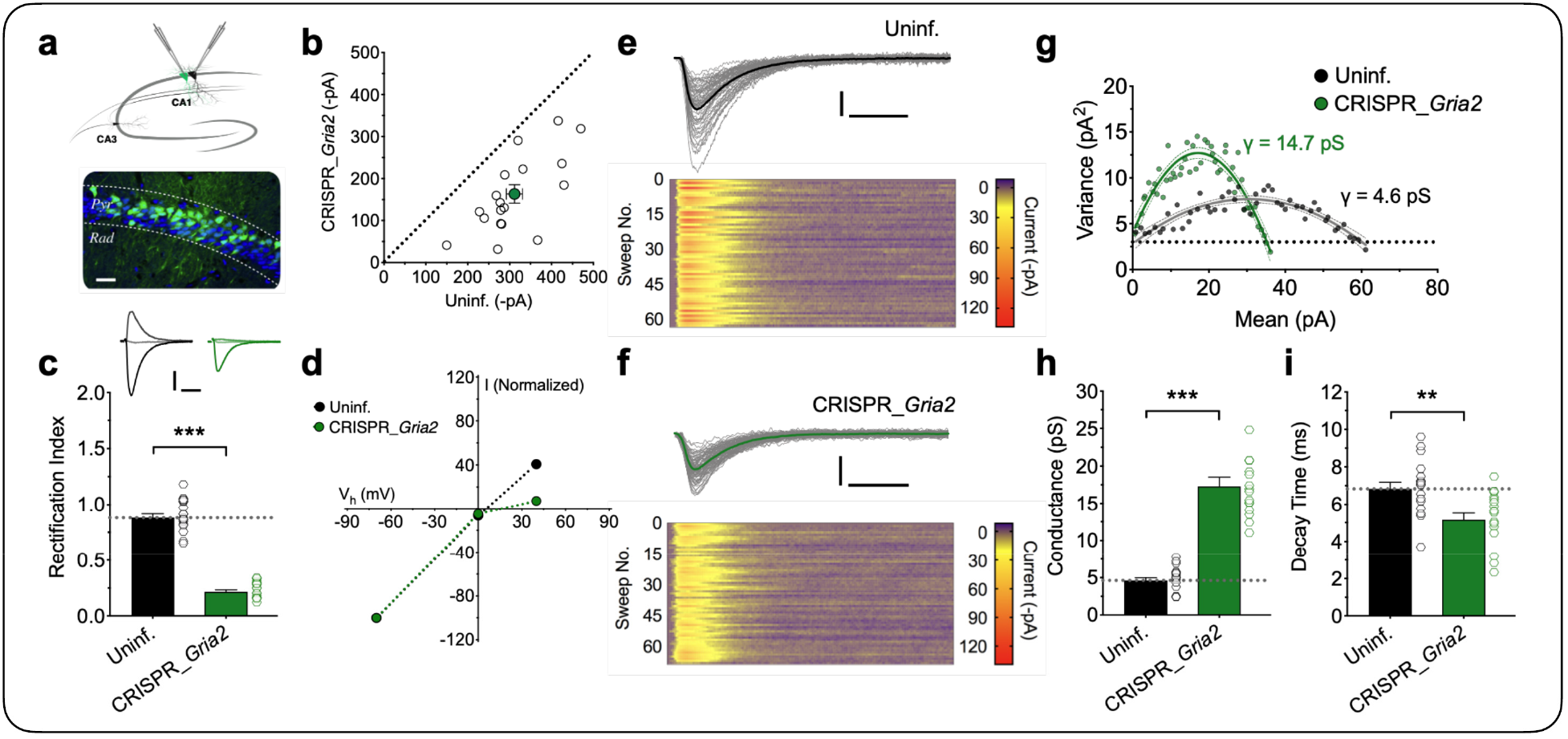
CP-AMPAR characterization in CRISPR_*Gria2* knock-out neurons. **a**, Schematic of dual whole-cell recordings for the CRISPR_*Gria2* knock-out and neighbouring uninfected (Uninf.) neurons. Sparse expression following stereotactic lentivirus injection detected by coexpressed EGFP (green); blue, DAPI staining; Pyr, stratum pyramidale; Rad, stratum radiatum. Scale bar = 30 microns. **b**, Scatterplot shows amplitudes of AMPAR EPSCs for each pair recorded simultaneously (open circles) and the mean ± SEM (filled circle; n = 18 pairs /12). **c-d**, Quantification of the rectification index for pharmacologically-isolated AMPAR-mediated EPSCs and the corresponding current-voltage relationship. Scale bars: 100 pA and 10 ms. **e-g**, Representative traces to measure the single channel γ for the control and CRISPR_*Gria2* knock-out neurons. Individual traces (thin lines) superimposed with the average. Scale bars: 30 pA and 10 ms. The lower panels are corresponding colour-coded images of all sweeps used in the NSFA (**g**). **h-i**, Quantification of γ and decay time constants (n = 17 and 16 for Uninf. vs. CRISPR_*Gria2* knock-out neurons from 12 animals).

## DISCUSSION

NMDA receptor-dependent LTP has been extensively studied as the primary mechanisms utilized are crucial for the formation of long-term memories. Despite many molecules being discovered and different aspects of their regulation being uncovered, there are crucial gaps in our knowledge. One relates to the fact that long-term memory requires *de novo* protein synthesis yet most of our mechanistic understanding of LTP has been obtained from the study of a protein synthesis-independent form of LTP. A second pertains to the fact that much of this understanding has been derived from the study of juvenile animals, where technical issues have permitted more in-depth analysis, whereas most studies of learning and memory are conducted in adult animals. In the present study, we have addressed these issues by studying LTP at CA1 synapses in young adult rodents and have compared induction protocols that are known to activate the protein synthesis-independent (cTBS) and protein synthesis-dependent (sTBS, rolipram + wTBS) forms ^35,36^. Using a cTBS protocol, LTP involved the insertion of additional CI-AMPARs, for which activation of CaMKII is both necessary and sufficient. Using a sTBS there was an additional LTP component that involved the transient insertion of CP-AMPARs, for which activation of CaMKII and PKA are both necessary and, in combination, sufficient. The insertion of CP-AMPARs increases AMPA receptor γ and this underlies the initial expression of this form of LTP, which we have termed LTP_γ_. The insertion of CP-AMPARs is transient and is replaced by a persistent increase in the number of CI-AMPARs.

### Two distinct postsynaptic forms of LTP at CA1 synapses

The division of NMDA receptor-dependent LTP into multiple components was made on the basis of sensitivity to various pharmacological agents and substantiated by genetic studies ^36^. In particular, when a single train (tetanus or TBS) is employed, the resultant LTP is independent of both PKA activation and *de novo* protein synthesis; this is commonly referred to as LTP1 or E-LTP ^35,37^. In contrast, when multiple trains are delivered, with an interval in the order of minutes, then there is the generation of an additional PKA and *de novo* protein synthesis-dependent component of LTP, which is commonly referred to as LTP2 or L-LTP ^27,36,38,39^. LTP2 is generally assumed to underlie long-term memory formation, that also requires *de novo* protein synthesis.

NMDA receptor-dependent LTP has also been divided into two distinct postsynaptic mechanisms of expression, one involving an increase in the number of AMPARs without a change in γ (LTP_N_) and the other involving an increase in γ (LTP_γ_) ^6^. Here, one of our goals was to determine whether these separate expression mechanisms specifically relate to LTP1 and LTP2. We found that LTP1 never involved an alteration in γ whereas LTP2 invariably did. The increase in γ was transient, lasting between 10 and 20 min and could be fully explained by the insertion of CP-AMPARs. In terms of signaling cascades, we found that activation of CaMKII was both necessary and sufficient for LTP1 whereas both CaMKII and PKA were required, and in combination were sufficient, for LTP2 (see model in Fig. 8). Our findings do not conflict with a large body of literature regarding alterations in AMPARs underlying LTP at these synapses and the roles of both CaMKII and PKA (e.g., ^26,30,40–42^). Hitherto, the roles of CP-AMPARs and alterations in γ in LTP have been controversial ^6,24,43–45^. However, these controversies can now be reconciled on the basis of the type of LTP under investigation.

**Fig. 8.**
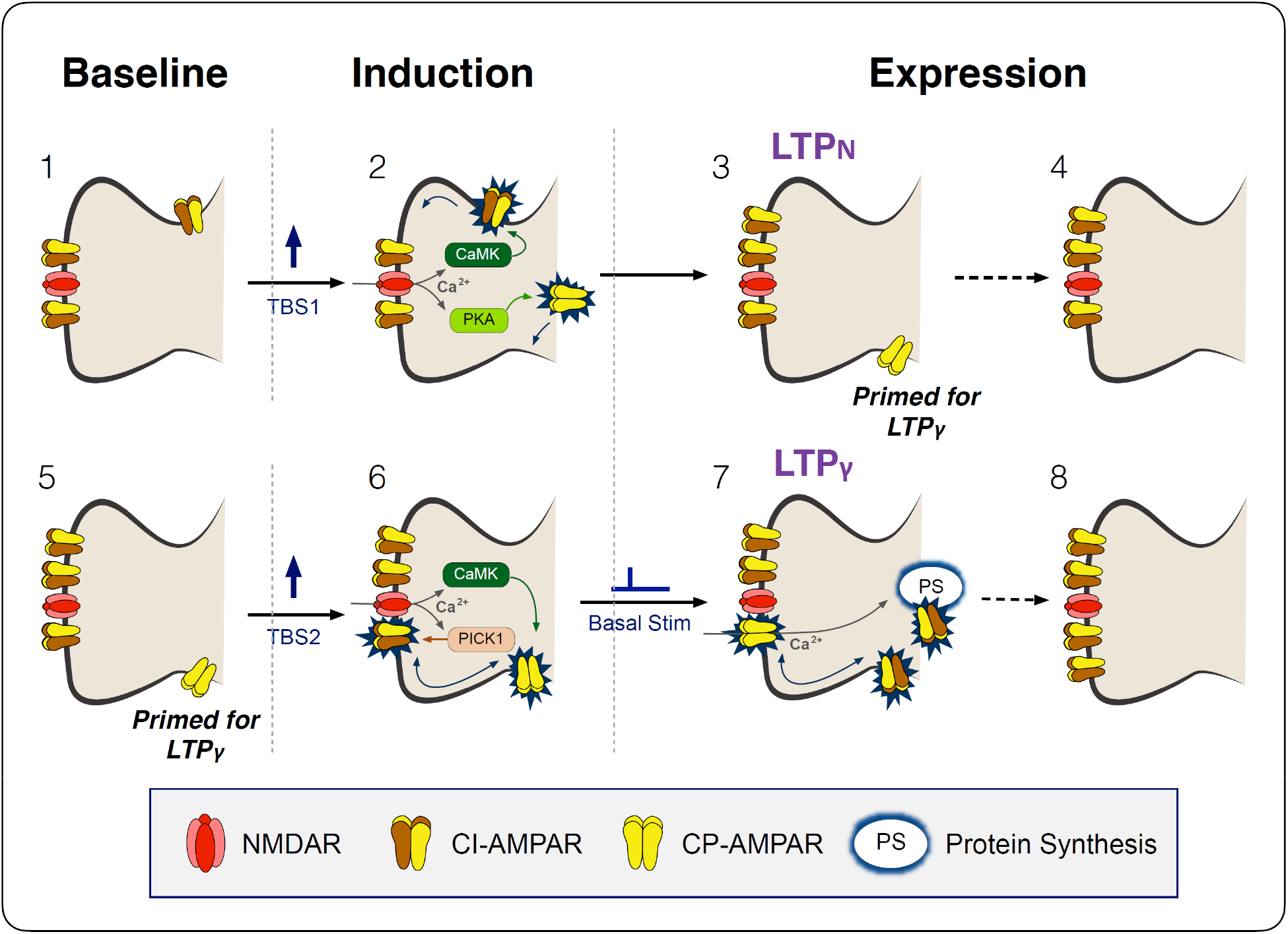
Schematic outlining the induction of two mechanistically distinct forms of LTP. **1.** Under baseline conditions synaptic transmission is mediated by GluA2-containing, calcium-impermeable (CI)-AMPARs, two shown for simplicity. **2.** The first theta-burst stimulation (TBS) activates NMDA receptors (NMDARs) and this drives more CI-AMPARs into the synapse by lateral diffusion from a peri-synaptic pool, via a process that involves CaMKII. PKA is also activated (via adenyl cyclase, not shown) and this induces the process of transducing GluA2-lacking calcium-permeable (CP)-AMPARs into peri-synaptic sites on the plasma membrane. **3.** LTP is expressed by the increase in number of CI-AMPARs (LTP_N_) but synapses also become primed for LTP_γ_ by the availability of peri-synaptic CP-AMPARs. **4.** Within ~ 1 h, the peri-synaptic CP-AMPARs are removed and, presumably, degraded. **6.** If a second TBS is delivered whilst the synapses are still primed (**5**) then NMDAR activation drives the peri-synaptically located CP-AMPARs into the synapse, via a CaMKII-dependent process. This might involve an exchange of CP-AMPARs for CI-AMPARs, which are removed from the synapse via a mechanism triggered by PICK1. **7**. These CP-AMPARs increase synaptic strength due to their higher single channel conductance (LTP_γ_). However, their dwell time in the synapses is quite short (~15 min) before they are removed. If synapses remain active, such as by basal stimulation, activation of the transiently-available, synaptic CP-AMPARs triggers protein synthesis and the insertion of more CI-AMPARs (**8**), which can extend the expression of LTP for long periods.

We can conclude therefore that LTP1 equates to LTP_N_ and LTP2 with LTP_γ_. It is important to note, however, that although a compressed induction protocol (cTBS) will ordinarily result in just LTP1 / LTP_N_ a spaced protocol will comprise a mixture of LTP1 and LTP2 (LTP_γ_), since the initial train will induce LTP1 upon which subsequent trains will add LTP2 under our experimental conditions. The relative proportion of these two components will depend on a variety of conditions, including the interval between the trains (with ~10 min being optimal for the induction of LTP2) and external factors such as stress, that likely impacts LTP1 and LTP2 differently ^31^.

### On the mechanism of LTP_γ_

Previous work has shown that an increase in AMPAR γ could result from either a CaMKII-dependent phosphorylation of Ser831 of GluA1 ^14,16^ or by the insertion of CP-AMPARs ^25^, since these have a higher single channel conductance than CI-AMPARs ^17,18^. Our findings have demonstrated that LTP_γ_ can be explained exclusively by the latter mechanism, since all changes in γ were eliminated by IEM. Furthermore, we found that activation of PKA plus CaMKII increased γ whereas CaMKII alone did not, despite leading to a substantial potentiation. The failure of CaMKII alone to increase γ, which is contrary to some previous studies ^14,15^, could be explained on the basis of the native AMPAR configuration since γ alterations are affected by the subunit combination and accessary protein composition of AMPA receptors ^16,18^. It is worth noting, however, that whilst activation of CaMKII alone was not sufficient to induce LTP_γ_ its activation was necessary. It is possible, therefore, that phosphorylation of Ser831 of GluA1 is necessary, but not sufficient for LTP_γ_.

Space precludes a detailed discussion of the underlying molecular mechanisms of LTP_γ_, but it is likely to be governed by AMPAR subunit-specific regulation and trafficking ^46^. In brief, our data are compatible with an exchange of a subset of CI-AMPARs for CP-AMPARs. The latter could be explained by a mechanism involving the Ca^2+^ sensor PICK1 ^47,48^, which has been shown to bind and internalize GluA2-containing AMPARs to enable the insertion of CP-AMPARs during LTP ^49^. The next step involves the replacement of the newly inserted CP-AMPARs with CI-AMPARs, a process that requires baseline (low frequency) synaptic activation ^26,43^ and probably involves Ca^2+^ permeation through the CP-AMPARs themselves ^50^. The rapid replacement of CP-AMPARs with CI-AMPARs was originally described at excitatory synapses onto cerebellar stellate neurons from P18-P20 rats ^51^. At this synapse, high frequency stimulation (tetanus) induces CP-AMPARs to be replaced with the equivalent number of CI forms resulting in a reduction in the synaptic current by a third, reflecting lower γ of the latter form. We observed an initial reduction in EPSC amplitude following the triggering of LTP, which might be explained, in part, by a one-to-one exchange of CP-AMPARs for CI-AMPARs. Additionally, the transient expression of CP-AMPARs could trigger an increase in the number of AMPAR slots at synapses that enables an increase in the number of CI-AMPARs above and beyond what can occur during LTP1.

Since CP-AMPARs increase synaptic conductance why does there need to be an exchange for a greater number of CI-AMPARs to maintain the enhanced synaptic response? One possibility is that the expression of CP-AMPARs at these synapses needs to be restricted in time due to potential excitotoxicity ^52^. Therefore, they can only provide a transient mechanism of expression whilst triggering the more persistent switch resulting in a larger number of CI-AMPARs.

### Developmental regulation of the expression mechanisms of LTP

There is strong evidence that the expression mechanisms of LTP are developmentally regulated. The co-existence of two mechanisms involving the insertion of CI-AMPARs and CP-AMPARs can account for the LTP at P14 ^6^ and in young adults, as observed herein. However, at around P7, LTP is associated with a decrease in γ ^22^, which is most likely explained by the replacement of CP-AMPARs with a larger number of CI-AMPARs. Early in development there is also the insertion of AMPARs into synapses that appear to lack AMPARs altogether; so-called “silent” synapses ^53,54^. A potential scenario is as follows: first synapses acquire CP-AMPARs, next these are replaced by more CI-AMPARs. Thereafter LTP can increase the number of these CI-AMPARs via two mechanisms, one of which involves the transient insertion of CP-AMPARs and one that does not.

There have been far fewer studies regarding the mechanisms of synaptic plasticity in tissues obtained from adult animals compared to juvenile animals, despite most learning and memory studies are conducted in adult animals. This is a concern when attempting to relate mechanisms of synaptic plasticity to learning and memory. Our present study, conducted exclusively in tissue from young adult animals, shows that two distinct forms of synaptic plasticity can be readily induced simply by altering the patterns of activation. Our result that a cTBS protocol induces LTP that does not involve an alteration in γ is consistent with another study in adult animals ^24^. Our finding that a sTBS induces an additional component of LTP that involves an increase in γ is the first evidence that such a process occurs beyond early developmental stages.

### Functional significance of two forms of LTP

This raises the question as to why there are two distinct mechanisms to increase the synaptic complement of CI-AMPARs. Previous work has shown that the insertion of CP-AMPARs is specifically associated with the PKA and protein synthesis component of LTP ^27^ It is reasonable to assume, therefore, that the transient insertion of CP-AMPARs is part of the machinery that triggers *de novo* protein synthesis and the consequential morphological changes (spine enlargement and/or new spine formation). In contrast, in the absence of *de novo* protein synthesis, the increase in synaptic CI-AMPAR number can support the increase in synaptic efficacy. Although both processes can increase synaptic strength lasting many hours *in vitro*, it seems probably that only the protein synthesis-dependent form triggers synaptic changes that underpin long-lasting memories (lasting from days to lifetimes). Indeed, it has been shown that spaced training with access to reward enhances the persistence of memory, and treatment with rolipram after training enhances memory retention ^55^. The requirement for PKA to trigger the protein synthesis-dependent form of LTP also provides the opportunity for extensive neuromodulation. Neurotransmitters, such as noradrenaline and dopamine, and stress hormones, such as corticosterone, may, via the insertion of CP-AMPARs, augment protein synthesisdependent LTP to enhance and/or prolong the persistence of the associated memory (e.g., ^30,31,42,56,57^).

### Concluding remarks

We have identified the molecular basis of two independent forms of LTP that co-exist at hippocampal synapses in young adult animals, the occurrence of which is controlled by the patterns of synaptic activation during induction. The existence of these two distinct LTP mechanisms goes a long way in explaining many of the controversies that have plagued the field. LTP1 can be induced by a cTBS and involves the insertion of CI-AMPARs, and for this to occur activation of CaMKII is both necessary and sufficient. A sTBS, however, triggers both LTP1 and LTP2. This latter form of LTP involves the transient insertion of CP-AMPARs and this requires activation of PKA in addition to CaMKII.

## METHODS

### Hippocampal slice preparation

Transverse hippocampal slices (400 μm) were prepared from male Sprague-Dawley rats (1-3 months of age). Animals were anesthetized with isoflurane and euthanised by decapitation in accordance with UK Animals (Scientific Procedures) Act of 1986. The brain was then removed and placed in ice-chilled slicing solution that contained (mM): 124 NaCl, 3 KCl, 26 NaHCO_3_, 1.25 NaH_2_PO_4_, 10 MgSO_4_, 10 D-glucose and 1 CaCl_2_, saturated with 95% O2 and 5% CO_2_. The hippocampi were rapidly isolated from the brain and sliced using a vibratome (Microslicer) while maintained in the slicing solution. The CA3 region was removed to suppress the upstream neuronal excitability, and the slices were transferred to an incubation chamber that contained the recording solution (artificial cerebrospinal fluid, ACSF; mM): 124 NaCl, 3 KCl, 26 NaHCO_3_, 1.25 NaH_2_PO_4_, 2 MgSO_4_, 10 D-glucose and 2 CaCl_2_ (carbonated with 95% O_2_ and 5% CO2). Slices were allowed to recover at 32-34°C for 30 min, and then maintained at 26-28 °C for a minimum of 1 h before recordings were made.

### Field excitatory postsynaptic potential (fEPSP) recordings

The extracellular electrophysiology was performed in both interface and submerged type chambers maintained at 32°C, and continuously perfused at 2-4 mL/min with oxygenated ACSF. The slope of evoked fEPSPs (V/s) was measured in the CA1 region of hippocampal slices and bipolar stimulating electrodes were used at a constant voltage intensity (0.1 ms pulse width) throughout the experiments. Signals were amplified using Axopatch 1D (Molecular Devices) and digitized with BNC-2110 (National Instruments) A/D board at a sampling rate of 20 kHz. Recordings were monitored and analyzed using WinLTP ^58^. Each specific experiment was conducted on a single slice from an animal, so the n-value reflects both the number of slices and animals used.

Two independent Schaffer collateral-commissural pathways (SCCPs) were stimulated alternately to obtain the evoked synaptic responses, each at a constant baseline frequency of between 0.033 to 0.1 Hz. Following a stable baseline period of at least 20 min, LTP was induced using thetaburst stimulation (TBS) delivered at the same basal stimulus intensity. An episode of TBS comprises 5 bursts at 5 Hz, with each burst composed of 5 pulses at 100 Hz. For LTP induced by compressed TBS (cTBS), three TBS episodes were delivered with an inter-episode interval (IEI) of 10 s. For spaced TBS (sTBS), the same number of episodes were given with an IEI of 10 min (see Fig. 1b). Representative sample traces are an average of 5 consecutive responses, collected from typical experiments (stimulus artefacts were blanked for clarity).

### Whole-cell patch clamp recording

Whole-cell recording was made with ACSF that contained 50 μM picrotoxin (Abcam) and 20 μM (+)-bicuculline (HelloBio) to prevent GABA_A_ receptor mediated contribution. CA1 pyramidal cells were visualized with IR-DIC optics (Zeiss). The whole-cell solution comprised (mM): 8 NaCl, 130 CsMeSO_3_, 10 HEPES, 0.5 EGTA, 4 Mg-ATP, 0.3 Na3-GTP, 5 QX-314 and 0.1 spermine. The pH was adjusted to 7.2-7.3 with CsOH and osmolarity was set to 285-290 mOsm/L. The peak amplitude of evoked EPSCs (pA) was monitored and analyzed using WinLTP ^58^. Two independent SCCPs were stimulated alternately, each at a baseline frequency of 0.1-0.5 Hz. Borosilicate glass pipettes were fire-polished with a final resistance of 2-4 MΩ. Access resistance (R_A_) was estimated by fitting whole-cell capacitance current with a double exponential, and experiments were only accepted for analysis if R_A_ varied by < 15%. R_A_ values were 8.8 ± 0.3 MΩ; range from 6.2-12.8 MΩ. Signals were amplified using an Axopatch 200B (Molecular Devices), filtered at 2-5 kHz, and digitized at 20 kHz using a BNC-2110 (National Instruments) A/D board.

Cells were voltage-clamped at −70 mV throughout unless otherwise indicated. LTP was induced using TBS delivered at basal stimulus intensity while in current-clamp mode, and was triggered within 10 min of whole-cell to prevent the dialysis effect. In some experiments, the PKA catalytic subunit (PKA Cα, 300 U/mL) and/or CaMKII (250 U/mL) were included in the internal solution. CaMKII was activated (1× NEBuffer for Protein Kinases; 50 mM Tris-HCl, 10 mM MgCl2, 0.1 mM EDTA, 2 mM DTT and 0.01% Brij 35; 200 μM ATP, 1.2 μM calmodulin and 2 mM CaCl2; incubated for 10 min at 30°C) or heat-inactivated (65°C for 20 min) as described in the supplier’s manual (New England Biolabs).

To ensure recording stability, extracellular field EPSPs were simultaneously monitored as described previously ^26^. Peak amplitude (pA) and initial slope (V/s) of EPSCs and fEPSPs were measured, and displayed on-line, using WinLTP ^58^. Whole-cell recordings were initiated following collection of at least 10 min of stable baseline assessed by extracellular recordings.

### Peak-scaled, non-stationary fluctuation analysis (NSFA)

The unitary conductance (γ) of AMPA receptors was estimated using NSFA according to ^6^ (see also ^19–21^). Whole-cell responses were carefully selected for analysis using WinWCP (University of Strathclyde, Glasgow) and Mini Analysis software on the basis of the following criteria: first, precise alignment of traces on the rise phase; second, no contamination by spontaneous or polysynaptic currents; third, complete decay from the peak EPSCs. The traces were analyzed and the variance of the decay was plotted as a function of the amplitude at that time point. The x-axis was divided into 50-bins of equal current decrement from the peak. The single channel conductance was estimated by fitting the plot to a second polynomial equation, *σ^2^* = *iI* - *I^2^N* + *b_1_*, where *σ^2^* is the variance, *I* is the mean current, *N* is the number of channels activated, *i* is the single channel current and *b_1_* is the background noise. In the conductance conversion (i.e. γ = i/V), the driving force (V) is the difference between the holding (−70 mV) and reversal potentials (assumed to be 0 mV).

The kinetics of the mean EPSC from each neuron was estimated in Clampfit (Molecular Devices) by measuring 20-80% rise time (τ_rise_) and the time constant for the decay (τ_decay_). Representative sample traces are the averages of all of the traces that were selected for analysis, superimposed with individual peak-scaled traces (10 successive sweeps), unless otherwise stated. Stimulus intensity was set to obtain a sporadic observation of transmission failures but high enough to obtain a reliable estimate of γ.

### Plasmid constructs and lentivirus production

The following oligonucleotide sequences were used to generate single guide RNA (sgRNA) for GluA2 knockout: forward (5’ to 3’) CACC G ctaacagcatacagataggt; reverse (5’ to 3’) AAAC acctatctgtatgctgttag C ^59^. These were annealed and ligated into the lentivirus backbone developed by the Zhang lab ^60^. The construct was modified and used with the CaMKIIα promoter for Cas9-P2A-EGFP expression.

Lentivirus was produced by transfecting Lenti-X 293T cells with pMD2.G, psPAX2 and lentiCRISPR ^60^. The 293T cells were maintained in serum-free UltraCULTURE media (supplemented with 4 mM L-glutamine, 2 mM GlutaMAX-I, 0.1 mM MEM non-essential amino acids, 1 mM sodium pyruvate, 1× penicillin/streptomycin). Three days after transfection, the supernatant was filter sterilized (0.45 μm pore membrane, Millipore) and ultracentrifuged at 20,000 rpm (Beckman Coulter) with an additional sucrose filtration. The lentivirus pellet was resuspended in Dulbecco’s PBS and kept at −80°C.

### *In vivo* stereotactic injections and dual whole-cell recordings

The surgical procedure was performed under sterile conditions in accordance with the Institutional Animal Care and Use Committee of Seoul National University. Male C57BL/6 mice (2-3 months of age) were anesthetized by intraperitoneal injection of a ketamine (130 mg/kg body weight) and xylazine (10 mg/kg) mixture. The anesthetized mice were immobilized on a stereotactic apparatus and the lentiviral medium (0.5 μL per each at a flow rate of 0.1 μL/min; 5 x 10^9^ TU/ml) was bilaterally injected at CA1 area using a microinjection syringe (Hamilton). The coordinates used were −1.7 mm posterior, ±1.2 mm lateral to bregma and −1.5 mm below the skull surface.

Following 4-6 weeks of expression, the hippocampal slices were prepared and whole-cell recordings were made as described above. EGFP-positive and neighbouring uninfected neurons were identified by epifluorescence microscopy and compared by dual whole-cell recordings. Rectification index was measured as described in ^26^. AMPAR currents were isolated using a mixture of D-AP5 (100 μM) and L-689,560 (5 μM). The index was calculated by taking the responses from −70, 0 and +40 mV of holding voltages. Following the recordings, brain slices were PFA-fixed, stained with DAPI, and imaged on a confocal microscope (Leica SP8).

### Compounds

Drugs were prepared as frozen stock solutions (stored below −20°C). Compounds were as follows: *N,N,H*,-trimethyl-5-[(tricyclo[3.3.1.1^3,7^]dec-1-ylmethyl)amino]-1-pentanaminiumbromide hydrobromide (IEM-1460; HelloBio); 4-(3-(cyclopentyloxy)-4-methoxyphenyl)pyrrolidin-2-one (rolipram; Abcam); 4-[(2*S*)-2-[(5-isoquinolinylsulfonyl)methylamino]-3-oxo-3-(4-phenyl-1-piperazinyl)propyl] phenyl isoquinolinesulfonic acid ester (KN-62; Tocris and HelloBio); D-AP5 (HelloBio); L-689,560 (Tocris); a catalytic subunit of protein kinase A (PKA Cα, New England Biolabs); Ca^2+^/calmodulin-dependent protein kinase II (CaMKII, New England Biolabs).

### Statistical Analysis

All treatment groups were interleaved with control experiments. Data are presented as mean ± SEM (standard error of the mean). Responses were normalised to the baseline prior to LTP induction unless otherwise stated. Statistical significance was assessed using (two-tailed) paired or unpaired Student’s *t*-tests or one-way ANOVA as appropriate; the level of significance is denoted on the figures as follows: \**p* < 0.05, \*\**p* < 0.01 and \*\*\**p* < 0.001.

## ACKNOWLEDGEMENTS

This work was supported by the MRC, ERC, CIHR Foundation Grant #154276 (G.L.C.) and Brain Canada Foundation (G.L.C.) through the Canada Brain Research Fund, with the financial support of Health Canada. G.L.C. is the holder of the Krembil Family Chair in Alzheimer’s Research. This work was also supported by The National Honor Scientist Program (NRF-2012R1A3A1050385) of Korea (B.-K.K.).

## AUTHOR CONTRIBUTIONS

P.P. and G.L.C. conceived the study. P.P. carried out the experimental work. K.-H.K. and J.-i.K. helped with the CRISPR experiments. T.M.S., Z.A.B., H.K., M.Z. and B.-K.K. provided advice and technical and/or financial support. P.P., J.G., C.A.B. and G.L.C. wrote the manuscript.

